# Evaluation of Topramezone on Zebrafish Retinoid Signaling

**DOI:** 10.1101/167296

**Authors:** Haixing Liu, Pengxing Xu, Yu Fan, Weizhi Zhang

## Abstract

Topramezone is a highly selective herbicide developed for broadleaf and grass weeds control in corn. In this study, the effects of topramezone on zebrafish, especially in retinoid signaling were investigated. Zebrafish embryos were treated with topramezone from 4 hours post-fertilization (hpf) to 144 hpf. Exposed to topramezone significantly reduced the retinal and retinoic levels compared to controls. The transcriptional expression levels of retinol dehydrogenase (*rdh1*), retinoic acid receptor subunit (*raraa*), retinal dehydrogenase (*raldh2*), retinol binding protein (*rbp1a*), and cellular retinoic acid binding protein (*crabp1a* and *crabp2a*) were significantly decreased. Our results suggested that topramezone significantly impaired zebrafish retinoid signaling during a short time exposure. However, treatment with topramezone significantly increased the mRNA expression levels of *zfblue*, *zfrho*, *zfgr1, zfuv*, and *zfred*. Our data demonstrated that topramezone treatment could interrupt retinoid signaling and further affect zebrafish eye development.

## INTRODUCTION

Topramezone [3-(4,5-dihydro-1,2-oxazol-3-yl)-4-mesyl-o-tolyl] (5-hydroxy-1-methylpyrazol-4-yl) methanone, which has been commercially introduced in 2006 [1-3], is a highly selective herbicide and widely used for broadleaf weeds and annual grass control in corn and wheat [3-7]. The aquatic environments have been polluted by herbicide due to surface runoff, direct overspray or drift while applying the herbicide [8-12].

Topramezone is a 4-HPPD inhibitor, which leads to increased serum tyrosine levels [1, 13-15]. Previous studies have reported that topramezone causes a dose-dependent elevated adverse effects on the thyroid levels in rats [16, 17]. Thus, topramezone has the potential risk to the wildlife and human health. Although topramezone has been widely used for 10 more years, few studies have investigated the effects of topramezone exposure to the aquatic organisms. Topramezone residues enter the aquatic ecosystems mainly due to the drift or runoff [18]. Thus, with the wide usage of the topramezone-based herbicides, it has raised to be a primary problem to evaluate the safety and environmental impacts of this pesticide in the environmental toxicology.

Retinoic acid (RA) acts in pattern formation and organogenesis during the vertebrate development [19-21]. The synthesis of RA from retinol need two sequential steps, first is the oxidation of retinol to retinal and then retinal is oxidized to RA [20, 22, 23]. Upon synthesis, RA can bind to the nuclear ligand-activated transcription factors, such as RA receptors-α, β and γ which can dimerize with RXRs-α, β and γ to regulate the target genes expression [24-26]. Abnormal RA signaling in vertebrate embryos impacts various organ developments, such as the branchial arches and the nervous system [27-30].

In this study, we aimed to access the effects of topramezone treatment on retinoid signaling and the zebrafish eye development. Zebrafish embryos were exposed to different concentrations of topramezone, the mRNA expression levels of the key genes involved in retinoid signaling pathway was examined.

## MATERIALS AND METHODS

### Chemicals

Topramezone, DMSO, standards for retinol, retinal and retinoic acid were purchased from Sigma Aldrich (St Louis, MO, USA). Chemicals used for retinoid measurement were of HPLC grade.

### Zebrafish maintenance and topramezone treatment

Wild-type zebrafish were maintained and raised as described previously [31, 32]. Developmental stages of zebrafish embryos were characterized as described previously [33].

Sixty zebrafish eggs were incubated in the topramezone exposure solutionn (0, 1, 10 and 100 mg/L) from 4 hpf to 144 hpf. Control group treated with 0.01% (v/v) DMSO. Water was changed twice daily. The embryos were collected, immediately frozen in liquid nitrogen and stored at −80°C for subsequent gene, protein and retinoid analysis at 144 hpf.

### RT-PCR

Twenty embryos from each treated group was collected and total RNA was extracted using TRIzol (Invitrogen) according to the manufacturer’s instruction, and single-stranded cDNA was systhesized as described previously [34, 35].

### Real-time qPCR

Real-time quantitative PCR (qPCR) was carried out as previous described [36, 37]. Relative gene transcription levels were determined after normalizing to the mRNA content of reference gene β-actin. The primers used in this study was described somewhere else.

### Retinoid measurement

The extraction and measurement of retinoid profiles was carried out as previously described [38].

### Statistical analyses

All data are presented as the mean ± SE and the statistical significance was set at p ≤ 0.05. All statistical analyses were performend using SPSS 18 software. Differences between two groups were evaluated by the one-way ANOVA test followed by Tukey’s test.

## RESULTS

### Exposed to topramezone caused developmental toxicity in zebrafish

Embryos were treated with 0, 1, 10 and 100 μg/L of topramezone from 4 hpf to 144 hpf, the survival rates of each treated group was 93 ± 2.2 %, 86 ± 3.1 %, 75 ± 6.3 % and 20 ± 2.4 %, respectively. These data showed a significant lower survival rates after 100 μg/L topramezone exposure. The larvae appeared developmental toxicity after 48 hpf, and the hatching rates were also significant decreased after 72 hpf in the higher topramezone treatment.

### Topramezone exposure affected the retinoid profiles

After treated with different concentration of topramezone, zebrafish larvae were collected and analyzed. Retinal contents were significantly decreased in the 10 μg/L topramezone treated embryos and RA levels were significantly lower in the topramezone-treated groups compared to the untreated controls (Table 1). The RA contents were reduced in a dose-dependent manner (Table 1).

**Table 1.** Retinoid profiles (ng/mg protein) in zebrafish embryos treated with topramezone

|  | 0 µg/L | 1 µg/L | 10 µg/L | 100 µg/L |
| --- | --- | --- | --- | --- |
| Retinol | 140.3 ± 11.2 | 135.4 ± 12.1 | 142.2 ± 5.8 | 132.2 ± 17.9 |
| Retinal | 90.1 ± 3.1 | 112.1 ± 3.4 | 70.8 ± 12.4 | 40.2 ± 10.5 |
| Retinoic acid | 16.2 ± 2.1 | 12.6 ± 2.1 | 6.8 ± 1.2* | N.D. ** |

N.D., not detected.
□ Significant differences compared to the control group (One-way ANOVA, followed by Tukey's test: $p < 0.05$ ).
□□ Significant differences compared to the control group (One-way ANOVA, followed by Tukey's test: $p < 0.01$ ).

### Topramezone treatment changed the mRNA expression levels

The transcriptional expression of key genes involved in intracellular retinol and retinal transport, such as *crbp1a*, were significant diminished (Fig. 1). The mRNA expression level of retinol dehydrogenase (*rdh1*), which converts retinol into retinal, was significantly decreased; however, the transcriptional expression of *raldh2*, which transforms retinal to RA, was significantly increased (Fig. 1). The transcriptional expression levels of *crabp1a* and *crabp2a*, which are the two isoforms of the cellular retinoic acid binding proteins, were significantly decreased in the embryos exposed to topramezone (Fig. 1). We also observed that the mRNA expression level of *raraa* was downregulated in a dose-dependent manner in the embryos treated with topramezone (Fig. 1). The mRNA expression levels of five key genes that encode the rhodopsin and ultraviolet, red, blue and green opsins, as *zfrho, zfuv, zfred, zfblue* and *zfgr1*, were increased in a dose-dependent manner in the topramezone exposed embryos (Fig. 2).

**Figure 1.**
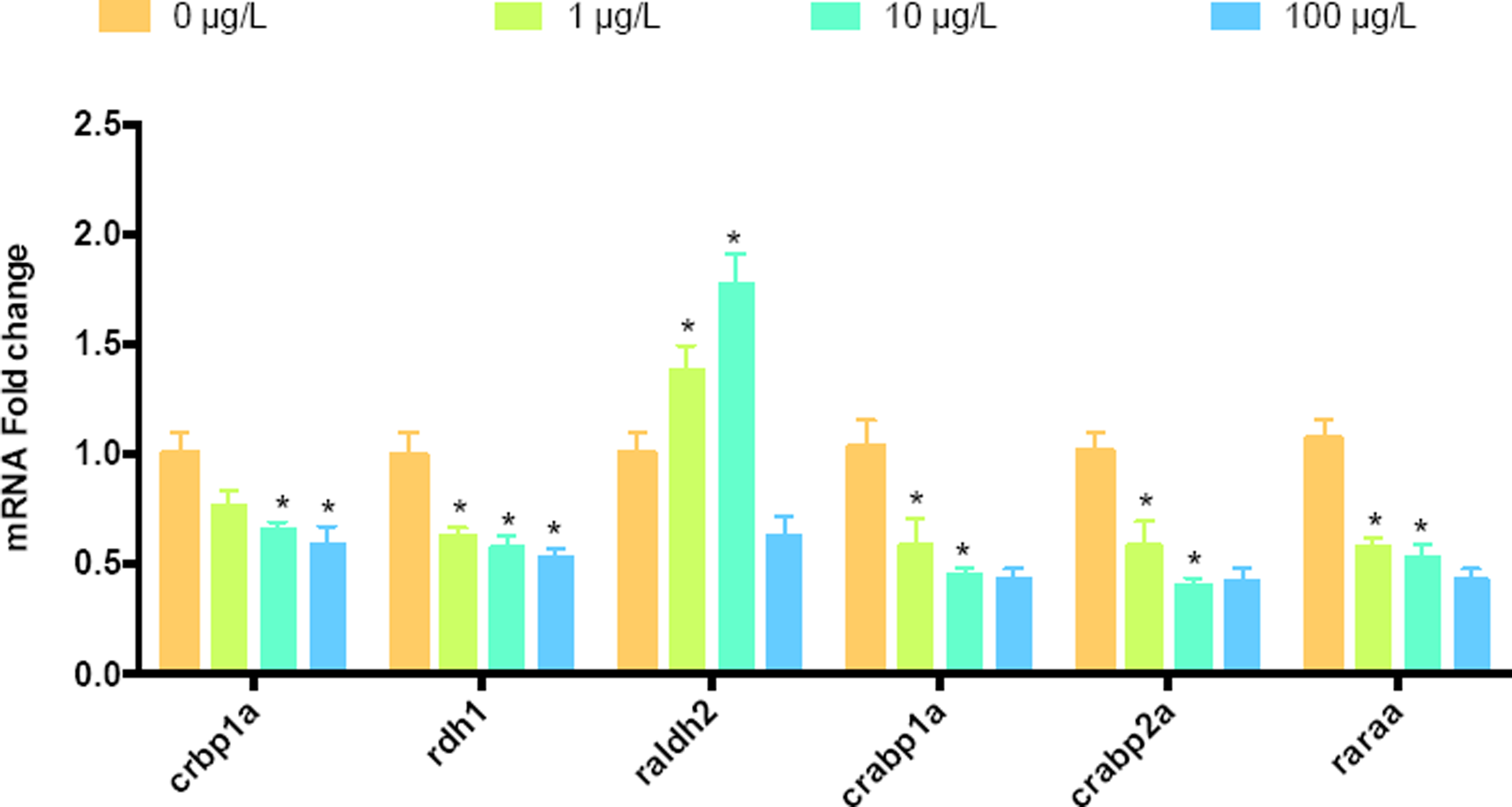
Transcriptional expression levels of *crbp1a, rdh1, raldh2, crabp1a, crabp2a* and *raraa* in zebrafish exposed to topramezone.

**Figure 2.**
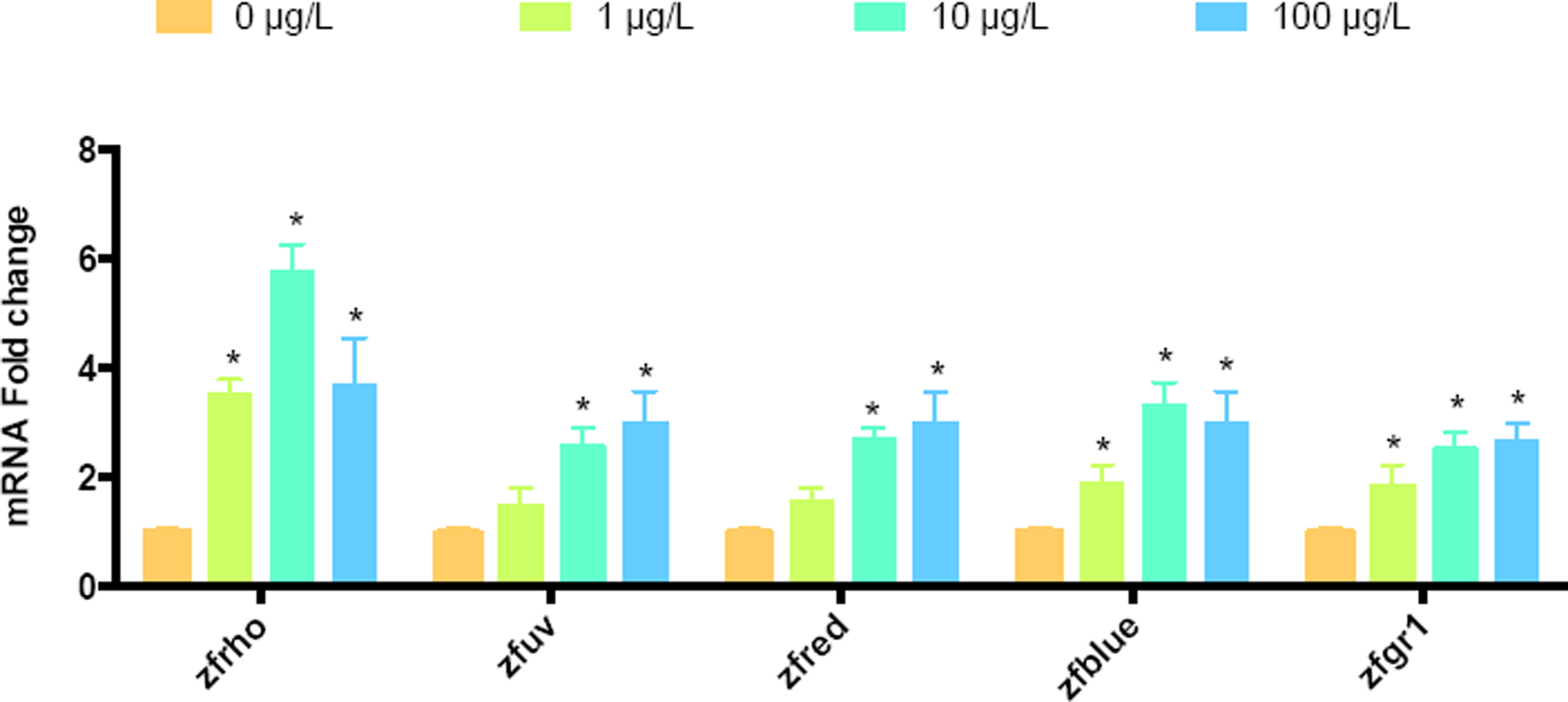
The mRNA expression levels of *zfrho, zfuv, zfred, zfblue* and *zfgr1* in zebrafish exposed to topramezone.

## DISCUSSION

The disruptive effects of topramezone treatment on retinoid contents and its effects on fish eye development are largely undefined. In the present study, we treated zebrafish embryos with topramezone to explore its role on the eye developmental process and related signaling pathway. Our results suggested that topramezone exposure could interrupt retinoid signaling and further affect the zebrafish eye development.

The total retinol levels had no significantly changes in the topramezone treated zebrafish larvae. Different responses to various concentrations of toxins might be explained by different exposed duration. Thus, treated with topramezone had no overt effect on the retinol contents in the zebrafish embryos might due to the short period of exposure time. Zebrafish eggs have high concentration of retinol [39], which indicates that those retinol preexisting in the embryos is enough for supporting the early stage development. When sugjected to the exogenous stress, those retinol can be mobilized and excreted into the plasma to maintain the developmental homeostasis [40].

The transcriptional expression of the retinol binding protein (*rbp4*) had no significant changes after treated with topramezone, suggesting that plasma retinol had been transmitted to target tissues normally. However, the contents of retinal were decreased in the higher concentration of topramezone exposure. These results suggested that the retinol cellular transport pathway was disrupted and caused little retinol being transformed into retinal.

RA has been reported to play an important role in regulating its production through a negative feedback mechanism by inhibition of raldh2 expression [41]. In the present study, the transcriptional expression of *raldh2* was up-regulated, which suggested that lower contents of RA could stimulate the production of RA. The mRNA expression level of *cyp26a* is independent of endogenous RA contents through *raldh2* in zebrafish embryos [42]. However, the transcriptional levels of *cyp26a* had no obviously changes, which indicated that there was a constant capability for RA degradation in zebrafish during embryonic development.

In zebrafish, *raraa*, partly functional overlaps with *raldh2* and *crabp2a*, that is existed in the hindbrain, tailbud and eye [43]. Thus, the downregulation of *raraa* mRNA expression levels in the present study demonstrated that the RA concentrations in target tissues were decreased.

RA plays a key role in the visual systems of vertebrates for photoreceptor development [44]. Many genes involved in eye morphogenesis are regulated by RA through its binding and interacting with the RARs and RXRs [45]. RA also plays a vital role in controlling the retina photoreceptors development in animals. Treatment of exogenous RA results in duplication of the retina during optic primordial development in fish [46]. Thus, the reduced RA contents might also had an adverse effect on the development of photoreceptors. In this study, the mRNA expression levels of *zfrho, zfuv, zfred, zfblue*, and *zfgr1*, that encode rhodopsin, ultraviolet, red, blue and green opsins, respectively, were significantly increased in the topramezone treated zebrafish larvae. These data indicated that reduced chromophore retinal and the disturbance of RA signaling in eye photoreceptors in the response to topramezone. Further studies are needed to evaluate the effects of long time exposure to relevantly environmental levels of topramezone on the retinoid homeostasis and eye development in the aquatic animals.

